# Translational control of AMPK activity in melanoma

**DOI:** 10.64898/2025.12.30.697000

**Authors:** Natália Vadovičová, Anna Lešková, Kateřina Koždoňová, Karolína Smolková, Barbora Valčíková, Filip Kafka, David Potěšil, Zbyněk Zdráhal, Ondřej Vacek, Ráchel Víchová, Karel Souček, Stjepan Uldrijan

## Abstract

The eIF4F translation initiation complex controls ERK MAPK signaling in malignant melanomas with *BRAF* and *NRAS* mutations. It also contributes to the development of melanoma resistance to therapies targeting BRAF and MEK kinases. Here, we uncovered a critical role for eIF4F in regulating the main cellular metabolic sensor, AMP-activated protein kinase (AMPK). In melanoma cells with the most common BRAF V600E mutation, ERK and AMPK pathway activities were reported as mutually exclusive. This is because BRAF-driven ERK activity negatively impacts LKB1-mediated canonical AMPK activation. However, we observed that eIF4F inhibition can stimulate AMPK activity in melanoma cells, both *in vitro* and *in vivo*, despite concomitant ERK hyperactivation. Notably, the protein levels of LKB1 and its co-factor MO25 were sensitive to eIF4F inhibition, indicating a non-canonical LKB1-independent mechanism of AMPK activation. In a proteomic screen, we aimed to identify eIF4F roles in melanoma cell physiology beyond the MAPK pathway. We found that the eIF4F function is essential for maintaining cellular levels of key cell cycle and metabolic regulators, including CDK1, CDK2, TYMS, and UHRF1. Importantly, we also identified the protein phosphatase PP2A as a new eIF4F target. Our subsequent analyses showed that inhibition or siRNA-mediated knockdown of PP2A increases AMPK activity in melanoma cells, independent of LKB1. This data shows that PP2A plays a significant role in regulating AMPK activity in melanoma. Thus, eIF4F inhibition not only impairs canonical AMPK activators but also downregulates PP2A, which negatively regulates AMPK dynamics.

Collectively, our data highlight a dual role of eIF4F in the control of AMPK in *BRAF*-mutant melanoma cells. It maintains the canonical AMPK signaling pathway while simultaneously limiting the extent of AMPK activation via the eIF4F-PP2A-AMPK axis. Pharmacological inhibition of this axis can overcome the negative control of AMPK signaling by the ERK pathway. This suggests new therapeutic opportunities to disrupt melanoma growth.

## Introduction

AMP-activated protein kinase (AMPK) is a key energy sensor and a context-dependent tumor suppressor that limits anabolic growth programs and promotes catabolic metabolism (1–4). Active AMPK is formed by three subunits: of α, β, and γ. Its activation depends on the phosphorylation of threonine 172 in the activation loop of the α-subunit by upstream kinases (5–8). The canonical activation mechanism of AMPK is mediated by liver kinase B1 (LKB1/STK11), which partners with STRAD and MO25 (CAB39) (9–12). LKB1 phosphorylates threonine 172 in response to cellular energy stress, which is marked by an elevated AMP:ATP ratio (5, 13). Increased AMP levels promote AMP binding to the regulatory γ-subunit of AMPK, inducing conformational changes that facilitate LKB1-mediated phosphorylation and protect the phosphorylated activation loop from dephosphorylation by protein phosphatases (7, 13–15).

In addition to the classical energy-sensing pathway mediated by LKB1, AMPK activation can occur through non-canonical, AMP-independent mechanisms (4, 16). These alternative routes include increases in intracellular calcium concentrations, sensing of glycolytic intermediates, and responses to DNA damage. The calcium-dependent pathway is mediated by CaMKKβ (calcium/calmodulin-dependent protein kinase kinase β), also known as CaMKK2, which phosphorylates AMPK at threonine 172 independently of LKB1 (17–20). Furthermore, CaMKKβ also appears to mediate AMPK activation in response to DNA-damaging agents and supports the survival of tumor cells under genotoxic stress (21). Another AMP/ADP-independent mechanism involved glucose starvation, which triggers AMPK activation by sensing the absence of the glycolytic intermediate fructose-1,6-bisphosphate (FBP) (22). When FBP is absent, aldolase promotes assembly of a lysosomal complex containing v-ATPase, Ragulator, axin, LKB1, and AMPK, thereby mediating AMPK activation. (22–24). Notably, both aldolase and the pentameric Ragulator protein complex also regulate the activity of the mechanistic target of rapamycin complex 1 (mTORC1), highlighting the lysosome as a shared regulatory hub for AMPK and mTORC1 (25–28).

Malignant melanoma is an aggressive cancer commonly driven by constitutive activation of the RAS/RAF/MEK/ERK mitogen-activated protein kinase (MAPK) pathway through mutations in the *BRAF* or *NRAS* genes (29, 30). In melanomas with the most common oncogenic mutation, BRAF V600E, overactivated ERK kinase and its downstream effector p90 ribosomal S6 kinase (RSK) can phosphorylate inhibitory sites on LKB1. This impairs LKB1-mediated AMPK activation (31). Conversely, active AMPK directly phosphorylates BRAF at Ser729, disrupting its association with Kinase Suppressor of Ras (KSR) scaffold proteins and dampening ERK pathway signaling (32). These reciprocal constraints have supported the view that strong oncogenic MAPK signaling in BRAF-mutant melanoma is intrinsically incompatible with AMPK activation.

Dysregulated activity of eukaryotic translation initiation factor 4F (eIF4F) complex can contribute to cancer development and resistance to therapy (33–35). In melanoma, enhanced eIF4F activity has been implicated in the development of resistance to clinical BRAF and MEK inhibitors (36). We recently discovered that eIF4F is not merely permissive for protein synthesis but also a crucial regulator of oncogenic MAPK signaling in melanoma (37). Sustained eIF4F activity maintains cellular protein levels of the dual-specificity phosphatase DUSP6, an essential negative regulator of ERK phosphorylation, thereby partially limiting oncogenic ERK signaling flux. Pharmacologic or genetic inhibition of eIF4F reduces DUSP6 abundance and promotes rapid ERK hyperactivation and increased expression of immediate-early genes such as *EGR1* and *FOS* (37).

In the current study, we aimed to gain new mechanistic insights into the role of eIF4F-dependent translation in the crosstalk between AMPK and ERK pathways in melanoma and to explain an unexpected observation: the simultaneous activation of the AMPK and MAPK pathways in *BRAF*-mutant melanoma cells treated with small-molecule eIF4F inhibitors. We found that eIF4F serves as a master regulator of AMPK in melanoma cells, controlling both canonical and non-canonical AMPK activation pathways. We describe a previously unrecognized eIF4F-PP2A-AMPK axis that appears crucial for limiting AMPK activity, thereby permitting melanoma growth under normal, unstressed conditions. Collectively, our findings may provide new opportunities for therapeutically targeting melanoma tumors resistant to current treatments.

## Results

### Inhibition of eIF4F promotes simultaneous activation of AMPK and ERK pathways in *BRAF*-mutant melanoma cells *in vitro* and *in vivo*

In our earlier work (37), we identified eIF4F as a central regulator of ERK pathway signaling flux in malignant melanoma. Its function proved critical to maintain normal cellular levels of the dual phosphatase DUSP6/MKP3, a major negative regulator of ERK in melanoma cells. Inhibition of eIF4F rapidly reduced DUSP6 expression and triggered ERK hyperactivation, which in turn strongly induced the transcription factors EGR1 and FOS (37). The current state of knowledge about AMPK regulation in malignant melanoma describes the oncogenic activation of the ERK pathway in cells bearing strongly activating *BRAF* mutations, e.g. BRAF V600E, as incompatible with the activation of the AMPK pathway. ERK and RSK, two kinases downstream of oncogenic BRAF, phosphorylate the canonical AMPK activator LKB1/STK11 kinase, preventing it from binding and activating AMPK (31). Conversely, when activated, AMPK phosphorylates BRAF at S729, disrupting its association with the adaptor protein KSR1, leading to the downregulation of the ERK signaling (32). AMPK activation by metabolic stress can block MEK activation by mutant BRAF in melanoma (38). Therefore, when analyzing the impact of a small molecule eIF4F inhibitor CR-1-31-B in an *in vivo* melanoma model based on the human BRAF-mutated A375 cell line, we observed with interest the simultaneous presence of a strong expression of the ERK target EGR1 and activated AMPK kinase, as evidenced by increased phosphorylation at threonine 172 (Thr172) of the AMPK α subunit, in lysates of tumors that responded to the drug by a pronounced decrease in DUSP6 levels (samples 712 and 714; Fig. 1A). Subsequent analyses in A375 melanoma cells cultured *in vitro* and treated with CR-1-31-B confirmed the unexpected concomitant activation of the ERK and AMPK pathways (Fig. 1B).

**1.**
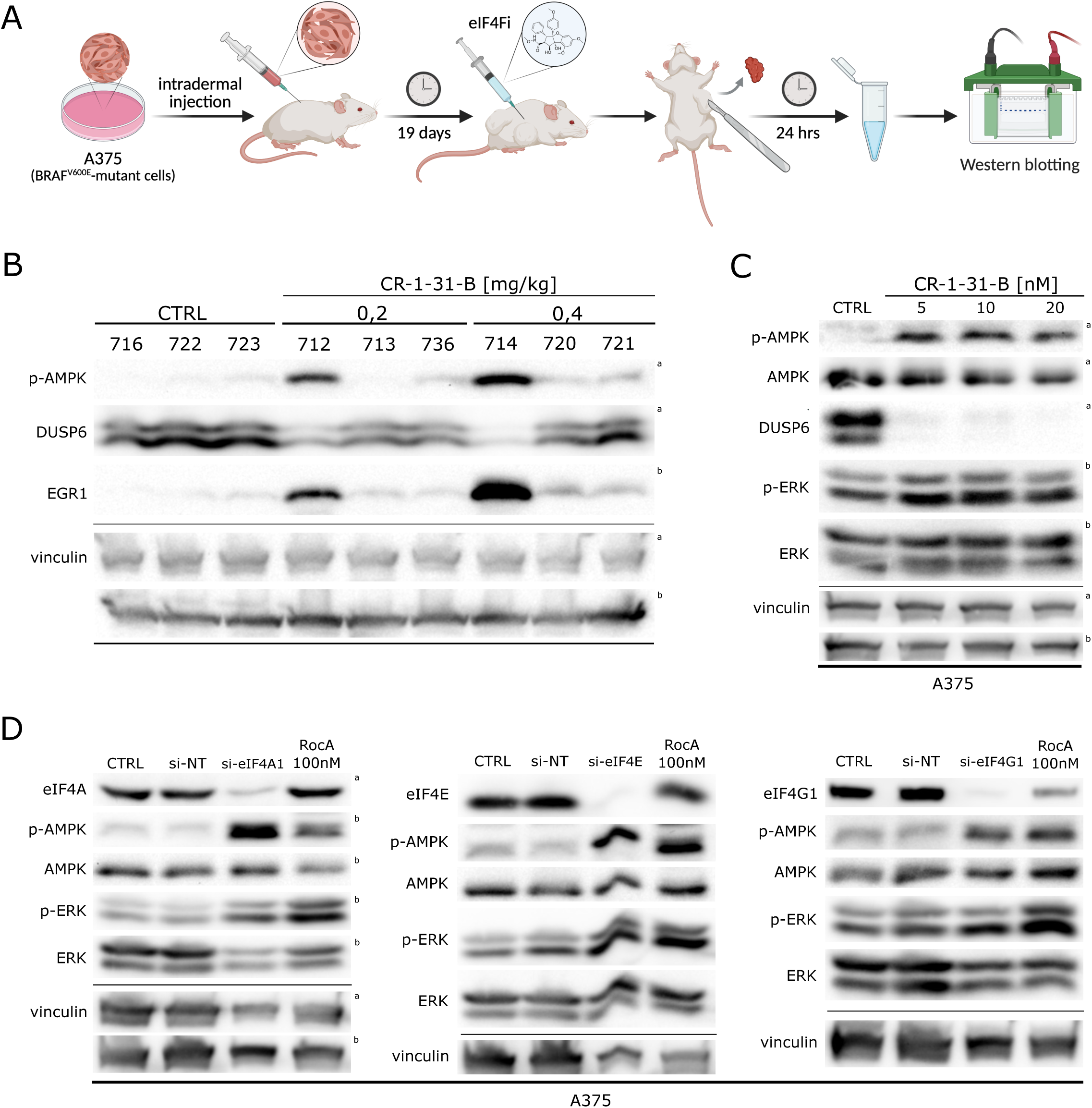
eIF4F inhibition promotes simultaneous ERK and AMPK activation both *in vivo* and *in vitro* in *BRAF*-mutant melanoma. **(A)** Schematic representation of tumor lysate preparation. A375 (BRAF^V600E-^mutant) melanoma cells were injected intradermally into mice. After 19 days, mice were treated with eIF4F inhibitor CR-1-31-B for 24 h. Tumors were collected, homogenized using the gentleMACS Dissociator system in lysis buffer containing protease and phosphatase inhibitors, and analyzed by western blot. Created with BioRender.com. **(B)** Western blot analysis of tumor lysates from A375 xenograft-bearing mice treated with the eIF4F inhibitor CR-1-31-B. DUSP6, EGR1, and phosphorylated AMPK (p-AMPKα Thr172) levels are shown. Tumors 712 and 714 exhibited marked DUSP6 depletion, elevated EGR1 expression, and increased AMPK phosphorylation. The control samples (CTRL) were treated with an equivalent volume of the vehicle (sesame oil). **(C)** Western blot analysis of A375 melanoma cells treated with CR-1-31-B for 20 h at the indicated concentrations. Treatment recapitulates the concurrent activation of ERK and AMPK pathways observed i*n vivo*. The control samples (CTRL) were treated with an equivalent volume of the vehicle (DMSO). **(D)** Western blot analysis of A375 cells following siRNA-mediated knockdown of eIF4F complex subunits (eIF4A1, eIF4E, and eIF4G1) for 48 h. Knockdown of each subunit decreased DUSP6 levels and increased phosphorylated AMPKα, confirming the eIF4F-specific mechanism of AMPK activation. Increased P-AMPKα was also observed upon treatment with Rocaglamide A (RocA, 100nM, 20 h), an eIF4F inhibitor. Non-targeting siRNA (si-NT) served as a control. The control samples (CTRL) were treated with an equivalent volume of the vehicle (DMSO). Vinculin served as a loading control. The upper index letters refer to the corresponding loading control detected on the same membrane.

To confirm that the observed AMPK activation was not due to an off-target effect of the selected small molecule compound, we performed gene knockdown experiments (Fig. 1C). We showed previously that targeted knockdown of genes encoding the eIF4F subunits eIF4A, eIF4E, and eIF4G via siRNAs promoted a drop in DUSP6 protein levels and subsequent ERK hyperactivation in human melanoma cells (37). In the current experiment, we also observed a significant increase in AMPK activity in A375 cells following the individual knock-down of genes coding for eIF4A1, eIF4E, and eIF4G1, supporting the eIF4F-specific mechanism (Fig. 1C).

### AMPK activation in melanoma by eIF4F inhibitors does not require LKB1

We next compared the impact of eIF4F inhibition on AMPK activity in the two most prevalent genetic subtypes of human melanomas. The A375 cell line, harboring the *BRAF* mutation, and the MelJuso line, carrying an *NRAS* gene mutation, were treated for 20 hours with rocaglamide A (RocA), a naturally occurring, highly selective inhibitor of the eIF4A RNA helicase subunit that is structurally related to the CR-1-31-B synthetic compound. As shown in Fig. 2A, both *BRAF*- and *NRAS*-mutated cells displayed a similar dose-dependent increase in AMPK activity following RocA treatment, revealing no subtype-specific difference. Unexpectedly, RocA also downregulated LKB1, the canonical AMPK-activating kinase (10, 12, 39), and its co-factor MO25/CAB39 (11), suggesting that eIF4F inhibition activates AMPK through an LKB1-independent pathway. The non-canonical AMPK activation was then confirmed using G361 human melanoma cells, which harbor the same BRAF V600E activating mutation as A375 but lack a functional *LKB1/STK11* gene. Even in the absence of LKB1, RocA treatment led to a dose-dependent AMPK activation and a decrease in MO25 levels (Fig. 2B). Next, we analyzed the melanoma response to a structurally unrelated eIF4F complex small molecule inhibitor 4E1RCat, acting by a molecular mechanism distinct from that of rocaglates CR-1-31-B and RocA. Importantly, all three tested human melanoma cell lines responded to 4E1RCat with AMPK activation, regardless of the status of the *LKB1/STK11* gene and the ERK pathway driver mutation (Fig. 2C).

**2.**
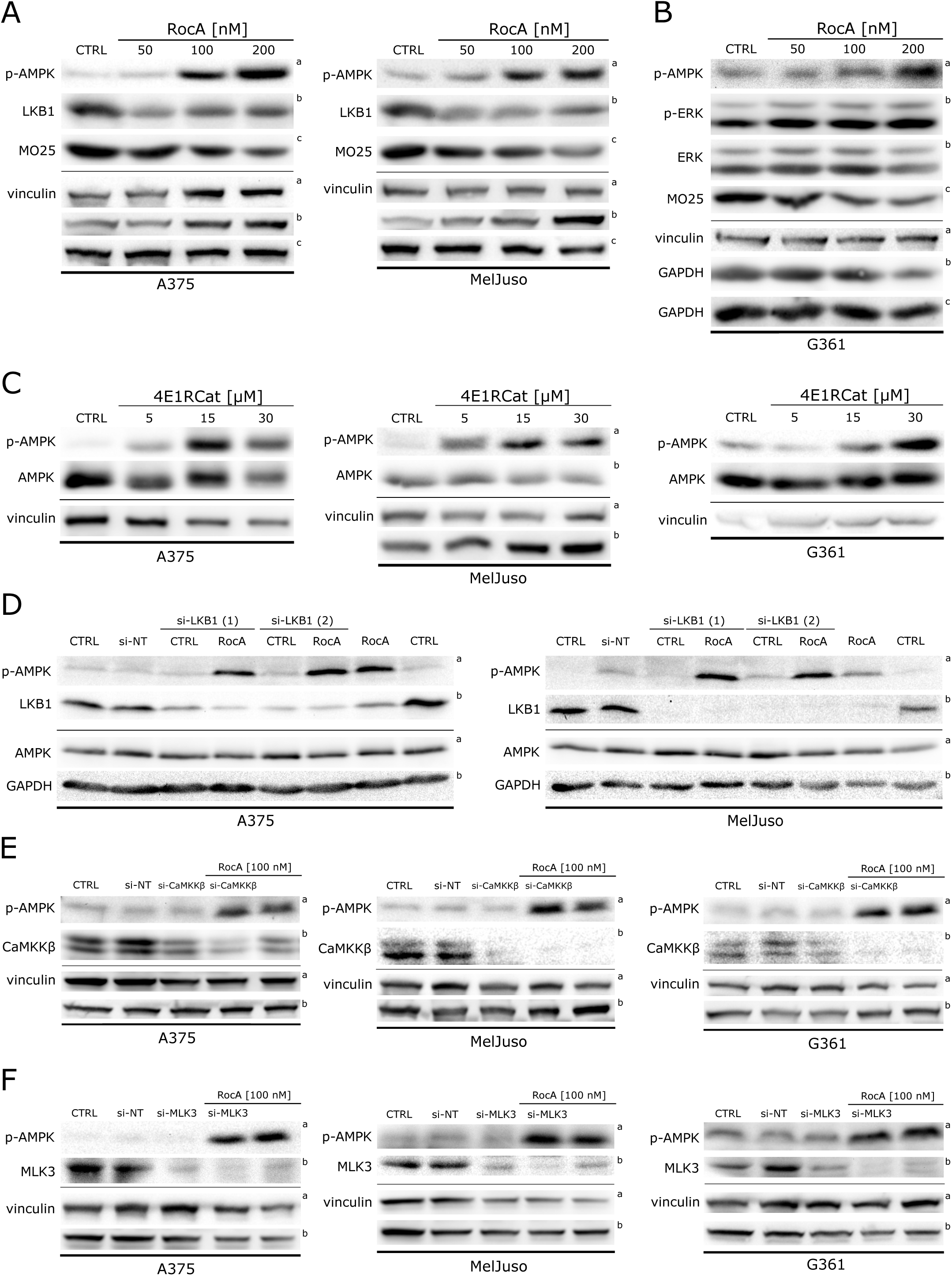
AMPK activation upon eIF4F inhibition in melanoma bypasses the canonical AMPK activator, LKB1. **(A)** Western blot analysis of A375 and MelJuso (NRAS-mutant) melanoma cells treated with increasing concentrations of the eIF4A inhibitor Rocaglamide A (RocA) for 20 h. Both cell lines exhibited dose-dependent increases in phosphorylated AMPK, accompanied by decreased levels of LKB1 and its cofactor MO25. **(B)** Western blot analysis of G361 (BRAF^V600E^-mutant, *LKB1*-null) melanoma cells treated with increasing concentrations of RocA for 20 h. AMPK activation occurred in a dose-dependent manner despite the absence of functional LKB1, confirming LKB1-independent AMPK activation. **(C)** Western blot analysis of A375, MelJuso, and G361 cells treated with increasing concentrations of the eIF4E-eIF4G disruptor 4E1RCat for 20 h. All three cell lines exhibited increased AMPK activity regardless of *LKB1* status or driver mutation. **(D)** Western blot analysis of A375 and MelJuso cells transfected with two different LKB1-targeting siRNAs for 48 h, combined with 100 nM RocA treatment for the last 20 h. LKB1 knockdown did not prevent RocA-induced AMPK activation, confirming the LKB1-independent mechanism. Non-targeting control siRNA (si-NT) was used as a control. **(D)** Western blot analysis of A375, G361, and MelJuso cells transfected with CaMKKβ-targeting siRNAs for 48 h, combined with 100 nM RocA treatment for the last 20 h. CaMKKβ depletion did not impair RocA-induced AMPK activation, indicating that CaMKKβ is not required for AMPK activation by eIF4F inhibition. Non-targeting control siRNAs (si-NT) were used as a control. **(E)** Western blot analysis of A375 and MelJuso cells transfected with MLK3-targeting siRNAs for 48 h, combined with 100 nM RocA treatment for the last 20 h. MLK3 protein levels decreased following both RocA treatment and siRNA-mediated knockdown, but MLK3 depletion did not prevent RocA-induced AMPK activation. Non-targeting siRNAs (si-NT) were used as a control. Vinculin, GAPDH, or total AMPK levels served as loading controls. The control samples (CTRL) were treated with an equivalent volume of the vehicle (DMSO). The upper index letters refer to the corresponding loading control detected on the same membrane.

To further verify the LKB1-independent mechanism of AMPK activation by eIF4F inhibition in cells expressing normal LKB1 protein levels, we performed a targeted knockdown of *LKB1* gene expression. A375 and MelJuso cells were transfected with non-targeting control siRNAs and two different *LKB1*-targeting siRNAs for 24 hours, followed by a 20 h treatment with 100 nM RocA. These results confirmed that the knockdown of *LKB1* expression did not hinder the AMPK activation induced by the eIF4F inhibitor RocA (Fig. 2D).

The Ca2+/Calmodulin-dependent protein kinase kinase β (CaMKKβ), also known as CaMKK2, is a well-established non-canonical activator of AMPK acting independently of LKB1 (17, 18). Therefore, we next tested the possibility that CaMKKβ could be responsible for AMPK activation in eIF4F inhibitor-treated melanoma cells. A375, MelJuso, and G361 cells were transfected with non-targeting control siRNAs and siRNAs targeting the expression of CaMKKβ for 24 h, followed by a 20 h treatment with 100 nM RocA. Both RocA treatment and CaMKKβ siRNAs significantly decreased CaMKKβ protein levels in all three cell lines, suggesting that CaMKKβ production in melanoma is eIF4F-dependent. However, the decrease in CaMKKβ expression had no discernible negative effect on AMPK Thr172 activating phosphorylation (Fig. 2E). Interestingly, another putative non-canonical AMPK activator Mixed lineage kinase 3 (MLK3, also known as MAP3K11) (40) also proved to be unstable in melanoma cells and requiring eIF4F activity for maintaining its cellular levels, and its knockdown had no significant impact on the AMPK activation induced by RocA (Fig. 2F).

### Lysosomal Ragulator complex is dispensable for AMPK and ERK modulation by high metabolic stress and eIF4F inhibition in *BRAF*-mutant melanoma

The finding that eIF4Fi-induced AMPK activation in melanoma cells was independent from LKB1 explained how it could happen despite the presence of the BRAF V600E mutation with its negative effect on LKB1 function (31). Nevertheless, the expected outcome of AMPK activation would be the downregulation of oncogenic BRAF-driven ERK signaling (32, 38), while eIF4F inhibition and knockdown promoted simultaneous activation of both AMPK and ERK signaling pathways. Moreover, in a recent study, we observed that a co-treatment with the AMPK activator AICAR could significantly potentiate eIF4Fi-induced ERK activity in *BRAF*-mutant melanoma cells (37). Therefore, we decided to have a more detailed look into the crosstalk of BRAF oncogene driven ERK signaling and AMPK activity to explain the unexpected concomitant activation of both signaling pathways in eIF4F-inhibited melanoma cells.

In our previous work, we reported a novel link between the metabolic state of melanoma cells and the activity of the oncogene-driven ERK pathway that was, however, very much dependent on the genomic context (38). We showed that in *NRAS*-mutant melanoma cells, low and high levels of metabolic stress potentiated oncogene-driven ERK signaling by promoting NRAS-independent interactions between KSR proteins and CRAF kinase. In contrast, in *BRAF*-mutant melanoma cells, the response of the ERK pathway to metabolic stress depended on the extent of AMPK activation. While low-level metabolic stress also further stimulated ERK activity by promoting the interactions between KSR and BRAF V600E proteins, high metabolic stress led to the recruitment of AMPK into the BRAF-KSR complex, followed by the disruption of the complex and inhibition of *BRAF* oncogene-driven ERK signaling (38). Based on these findings, we hypothesized that AMPK activity induced in *BRAF*-mutant cells in response to eIF4F inhibition might correspond to that induced by the low-level metabolic stress, still not reaching the threshold leading to the disruption of KSR-BRAF interaction and ERK inhibition. To test this possibility, we manipulated the metabolic state of BRAF V600E melanoma cells and compared the impact of eIF4Fi and low and high metabolic stress on AMPK activity and ERK signaling. We treated BRAF V600E melanoma cell lines A375 and G361 for 14 hours with a small molecule hexokinase inhibitor 2-deoxyglucose (2DG) to induce a low-level metabolic stress and mild AMPK activation, while treatments with 2DG in combination with the mitochondrial complex I inhibitor rotenone (Rot) were used to induce high levels of metabolic stress yielding strong AMPK activation.

As shown in Fig. 3A, the levels of active AMPK induced by RocA in *BRAF*-mutant melanoma cells were similar to those induced by 2DG, significantly lower than levels corresponding to strong AMPK activation observed in response to high metabolic stress (2DG + Rot). These data could explain the observed differential impact of the treatments on ERK activity. While high levels of cellular AMPK activity induced by the metabolic stressors blocked the ERK signaling, presumably by disrupting the BRAF-KSR association, a comparatively lower AMPK activity induced by eIF4Fi likely did not reach the threshold to disrupt the upstream ERK signaling, allowing for the amplification of the signal at the level of ERK kinase due to the disrupted negative feedback. To our surprise, however, the response of the LKB1-negative G361 cells essentially replicated the results obtained with LKB1-proficient A375 cells, suggesting that the potent upregulation of AMPK and ERK pathways by high metabolic stress in *BRAF*-mutant melanoma may not require LKB1 function.

**3.**
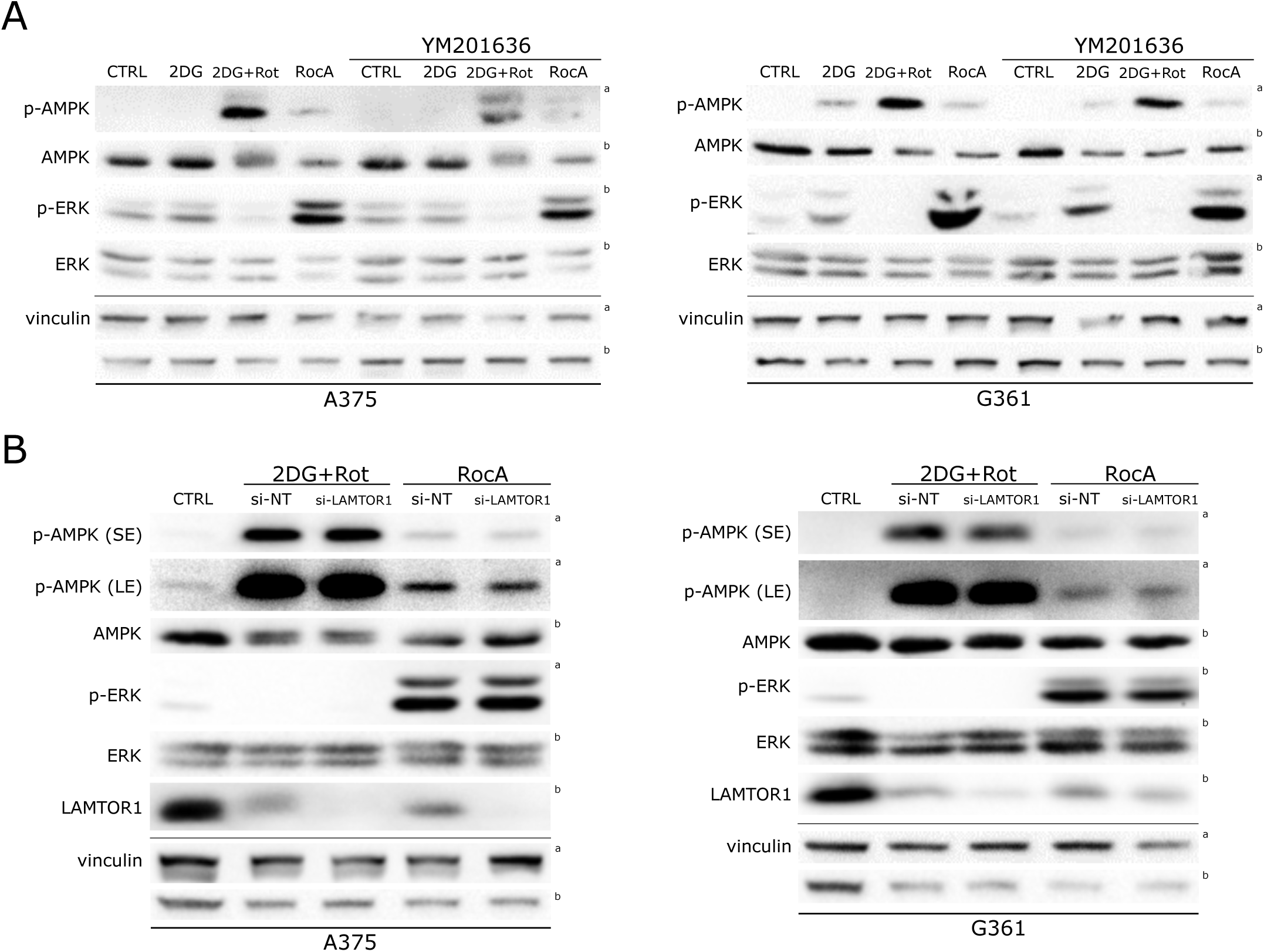
Non-canonical AMPK activation upon eIF4F inhibition and metabolic stress occurs independently of Ragulator complex in BRAF^V600E^ melanoma. **(A)** Western blot analysis of A375 and G361 melanoma cells treated with RocA (100 nM, 24 h), 2-deoxyglucose (2DG, 5,5 mM, 14 h), a combination of 5,5 mM 2DG and 5 µM rotenone (Rot), and 1 µM YM201636 (24 h) in combination with metabolic stressors or RocA, which were added for the last 14 h or 20 h of treatment, respectively. RocA and 2DG induced comparable low-level AMPK activation and maintained ERK activity, while high metabolic stress (2DG + Rot) triggered strong AMPK activation and ERK inhibition in both cell lines, regardless of LKB1 status. YM201636 co-treatment had no significant impact on AMPK or ERK activity, indicating Ragulator-independent regulation. Non-targeting control siRNAs (si-NT) were used as a control. **(B)** Western blot analysis of A375 and G361 cells transfected with LAMTOR1-targeting siRNAs for 48 h, combined with 100 nM RocA or 5,5 mM 2DG + 5 µM rotenone (Rot) treatment for the last 20 or 14 h, respectively. LAMTOR1 knockdown did not prevent the differential regulation of AMPK and ERK by eIF4F inhibition versus high metabolic stress, confirming Ragulator-independent mechanisms. Both RocA and 2DG + Rot treatments decreased LAMTOR1 protein levels. Non-targeting siRNAs (si-NT) were used as a control. The control samples (CTRL) were treated with an equivalent volume of the vehicle (DMSO). Vinculin served as a loading control. p-AMPK levels are shown with short (SE) and long (LE) membrane exposure times. The upper index letters refer to the corresponding loading control detected on the same membrane.

Different levels of metabolic stress seem to activate distinct AMPK pools in the cytoplasm and various cellular organelles (24). Intriguingly, the AMPK and ERK signaling pathways were reported to intersect on the lysosome. Specifically, in a protein complex called Ragulator that is anchored on the lysosome by its LAMTOR1/p18 subunit and is best known for its vital role in the activation of the mammalian target of rapamycin (mTOR) signaling pathway (26, 41–47). To disrupt the lysosomal compartment and the Ragulator complex function, we used YM201636, a small molecule inhibitor of the phosphoinositide kinase PIKfyve (48), which can interfere with later stages of endocytosis in melanoma cells, leading to the accumulation of enlarged lysosomes with LAMTOR1 sequestrated at their interface (49). The cotreatment with YM201636 did not have significant impact on either AMPK or ERK activity, suggesting that the Ragulator complex may not be involved in the crosstalk of the studied signaling pathways in melanoma in response to metabolic stress and eIF4F inhibition (Fig. 3A).

To test this possibility further, we performed siRNA-mediated knockdown of *LAMTOR1* in A375 and G361 cells and analyzed its impact on ERK and AMPK activity in the context of RocA and 2DG+Rot treatments. Data shown in Fig. 3B confirm the differential impact of high metabolic stress and eIF4F inhibition on ERK and AMPK activity and support the independence of their regulation on the Ragulator complex. It is further supported by the observation that eIF4Fi and high metabolic stress significantly downregulated LAMTOR1 protein levels in melanoma cells, both treatments possibly through their negative effect on protein synthesis. The highly similar response of A375 and LKB1-negative G361 cells to the treatments again suggested that non-canonical AMPK activation in malignant melanomas with BRAF V600E mutation may take place under high metabolic stress (Fig. 3B).

### Proteomic analyses identified dependency of key cell cycle and DNA replication regulators on continuous eIF4F activity in melanoma cells

Next, we employed a proteomic approach to uncover novel targets within the eIF4F pathway that could be linked to AMPK activation and the antitumor effects of eIF4F inhibitors in melanoma in general (Fig. 4A). We treated A375 and MelJuso melanoma cells with RocA (100 nM) for 24 hours. Control cells were treated with an equivalent amount of DMSO. Proteomic LC-MS analysis revealed a potential downregulation of 131 proteins in A375 cells and 201 proteins in MelJuso cells following RocA treatment (Suppl. Dataset S1). Fifty-seven proteins were identified as potential eIF4F targets in both melanoma models. Crucially, consistent with the results of our western blot analyses, the proteomic screen identified MO25/CAB39, the co-factor for the canonical LKB1-mediated AMPK activation, as one of the 57 proteins significantly downregulated in response to eIF4Fi in both melanoma cell lines. Among potential eIF4F downstream targets there were several cellular metabolic enzymes, e.g., ATP citrate synthase (ACLY), ornithine aminotransferase (OAT), and fatty acid synthase (FASN), indicating a potential impact of eIF4F inhibition on cancer cell metabolism. The list of identified eIF4F targets also contained essential cell cycle regulators, including the cyclin-dependent kinases CDK1, CDK2, and CDK6. In addition, levels of essential proteins participating in DNA replication and repair, thymidylate synthase (TYMS) and proliferating cell nuclear antigen (PCNA), were also identified by the proteomic analysis as sensitive to eIF4F inhibition in both melanoma subtypes. The screen results were also analyzed using Metascape (https://metascape.org) to cluster the identified proteins based on their known biological function and pathways (50). Among most prominent were functional clusters linked to cell cycle regulation, receptor tyrosine kinase signaling, microtubule cytoskeleton organization, membrane trafficking, DNA metabolism, PKR signaling, and antiviral responses (Suppl. Dataset S2).

**4.**
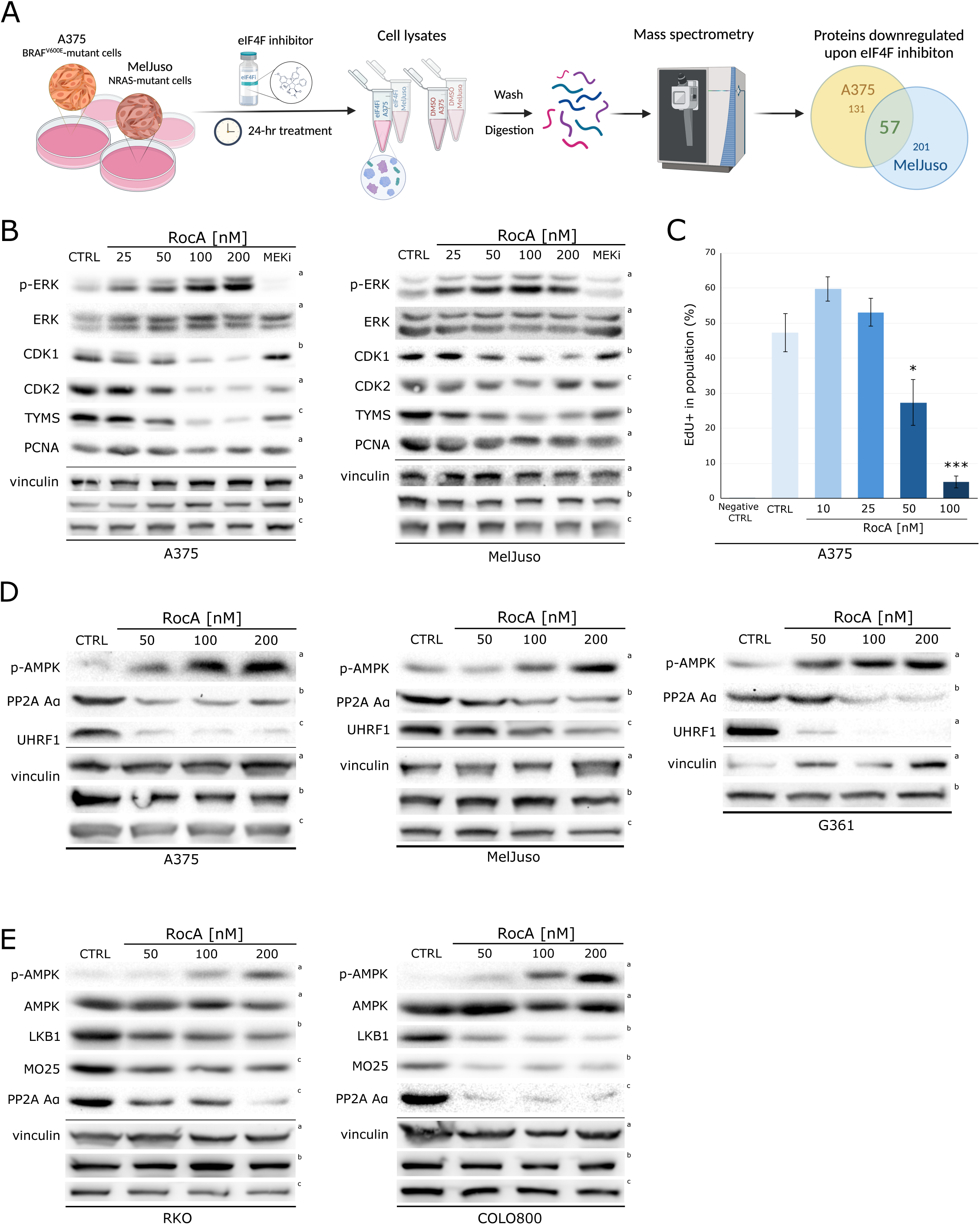
eIF4F activity maintains expression of key cell cycle regulators and AMPK negative regulators in melanoma. **(A)** Schematic overview of the mass spectrometry-based proteomic analysis of eIF4F inhibition in A375 and MelJuso melanoma cells treated with 100 nM RocA or DMSO vehicle control for 20 h. Created with BioRender.com. **(B)** Western blot analysis of A375 and MelJuso cells treated with increasing concentrations of RocA for 20 h. MEK inhibitor PD184352 (MEKi, 1µM) was used as a positive control for MAPK pathway inhibition. eIF4F inhibition resulted in dose-dependent decreases in CDK1, CDK2, TYMS, and PCNA protein levels, accompanied by ERK hyperactivation (p-ERK). **(C)** Flow cytometry analysis of DNA synthesis in A375 cells using Click-iT® EdU incorporation assay. Cells were treated with increasing concentrations of RocA for 18 h, followed by a 2 h incubation in fresh medium with 10 µM EdU added for the final 30 min. RocA concentrations ≥50 nM significantly reduced the proportion of EdU-positive cells. The control samples (CTRL and Negative CTRL) were treated with an equivalent volume of the vehicle (DMSO), and the negative control (Negative CTRL) was not pulse-labeled with EdU. Results were obtained from three independent repetitions, and the data are presented as the average percentage of EdU-positive cells± SE; p < 0.05. **(D)** Western blot analysis of A375, MelJuso, and G361 cells treated with increasing concentrations of RocA for 20 h. eIF4F inhibition led to dose-dependent decreases in UHRF1 and PP2A Aα subunit protein levels, accompanied by AMPK activation in all three cell lines. **(E)** Western blot analysis of COLO800 (BRAF^V600E^-mutant melanoma) and RKO (BRAF^V600E^-mutant colorectal cancer) cells treated with increasing concentrations of RocA for 20 h. eIF4F inhibition decreased LKB1, MO25, and PP2A-Aα subunit levels while activating AMPK, demonstrating that eIF4F-mediated control of AMPK activity extends beyond melanoma to other BRAF-mutant cancers. Vinculin served as a loading control. The control samples (CTRL) were treated with an equivalent volume of the vehicle (DMSO). The upper index letters refer to the corresponding loading control detected on the same membrane.

Next, we performed validation of selected potential targets connected to DNA synthesis and cell cycle progression using western blotting. The results confirmed that RocA-mediated eIF4F inhibition for 20 hours resulted in a marked concentration-dependent decrease in the expression levels of CDK1, CDK2, and TYMS in both melanoma subtypes, along with the expected hyperactivation of the ERK pathway, that served as a positive control (Fig. 4B). Interestingly, CDK2 and TYMS may represent dual eIF4F and ERK pathway targets as their levels in A375 melanoma cells decreased also in response to MEK inhibition. The reduction in PCNA levels was moderate, suggesting that this protein has a longer half-life compared to CDKs and TYMS, and is therefore less sensitive to the inhibitory effect of RocA on protein synthesis. These findings indicated that eIF4F inhibitors may impact DNA synthesis and the cell cycle progression in melanoma cells through downregulation of specific proteins. To study this possibility, using Click-iT™ EdU Alexa Fluor™ 488 Flow Cytometry Assay Kit, we measured the rate of the incorporation into newly synthesized DNA of EdU (5-ethynyl-2’-deoxyuridine), a nucleoside analog to thymidine. A375 melanoma cells were treated with RocA for 18 hours, followed by a two-hour incubation with a fresh cell culture medium with EdU addition for the last 30 minutes before collecting the samples and flow-cytometric analyses. The proportion of cells performing DNA synthesis was decreased significantly in cells treated with RocA at 50 nM or above, while lower RocA concentrations had no negative effect (Fig. 4C), consistent with the observed changes of the studied eIF4F targets (Fig. 4B). Interestingly, cells treated with low RocA concentrations (10 and 25 nM) tended toward increased rate of DNA synthesis (Fig. 4C), possibly reflecting already enhanced ERK pathway activity. Taken together, these findings suggest that eIF4F might play a pivotal role in sustaining the expression of proteins essential for melanoma cell proliferation.

### Identification of the eIF4F-PP2A axis controlling AMPK activity in malignant melanoma

Given the negative effect of eIF4F inhibition on the levels of AMPK-activating kinases, we hypothesized that the observed increase in AMPK activity following eIF4F inhibition could also result from a disrupted negative feedback control rather than upstream kinase overactivation, in a manner similar to the disrupted control of oncogenic ERK signaling in melanoma, where eIF4F inhibitors induced potent ERK overactivation by rapidly disrupting the essential negative feedback mediated by the DUSP6 phosphatase (37). Intriguingly, our proteomic analyses indicated a potential disruption of Protein phosphatase 2A (PP2A) function upon eIF4F inhibition, which could have a positive impact on AMPK activity in melanoma cells as PP2A has been implicated in the negative control of AMPK by dephosphorylating the critical T172 residue of the AMPK α subunit (51, 52). In our screen (Suppl. Dataset S1), we identified two PP2A activity regulators as potential eIF4F targets – the scaffolding subunit Aα of the PP2A enzyme complex, encoded by the *PPP2R1A* gene, along with the ubiquitin ligase UHRF1 (Ubiquitin like with PHD and ring finger domains 1). While UHRF1 serves as an essential epigenetic regulator participating in the maintenance of DNA methylation, it was also reported to promote the interaction between AMPK and PP2A (53).

A western blot validation of the proteomic data in A375, MelJuso, and G361 cells confirmed that RocA treatment led to a concentration-dependent decrease in UHRF1 and PP2A Aα proteins in both melanoma genetic subtypes (Fig. 4D). Intriguingly, the impact of eIF4Fi on LKB1, MO25, and PP2A Aα protein levels and concomitant AMPK activation was recapitulated in a subsequent experiment not only in another *BRAF*-mutant human melanoma cell line COLO800 but also in human colorectal cancer cell line RKO driven by the BRAF V600E oncogenic mutation (Fig. 4E), suggesting that eIF4F may contribute to the control of AMPK activity also in other types of cancer harboring *BRAF* oncogenic mutations.

To further explore the role of PP2A in negatively regulating AMPK activity in melanoma, we used two structurally diverse small-molecule PP2A inhibitors, okadaic acid and a clinical drug candidate LB-100. Both inhibitors induced a significant AMPK activation in all three tested melanoma cell lines, strongly suggesting that PP2A contributes to the control of AMPK activity (Fig. 5A,B). However, to rule out any potential off-target effects on melanoma cells of the two small molecule compounds, we knocked down the PP2A Aα subunit expression using specific siRNAs, which led to a significant increase in active AMPK levels similar to that induced by RocA, confirming the vital role of the eIF4F-PP2A axis in limiting AMPK activity in human melanoma cells in the absence of metabolic stress (Fig. 5C).

**5.**
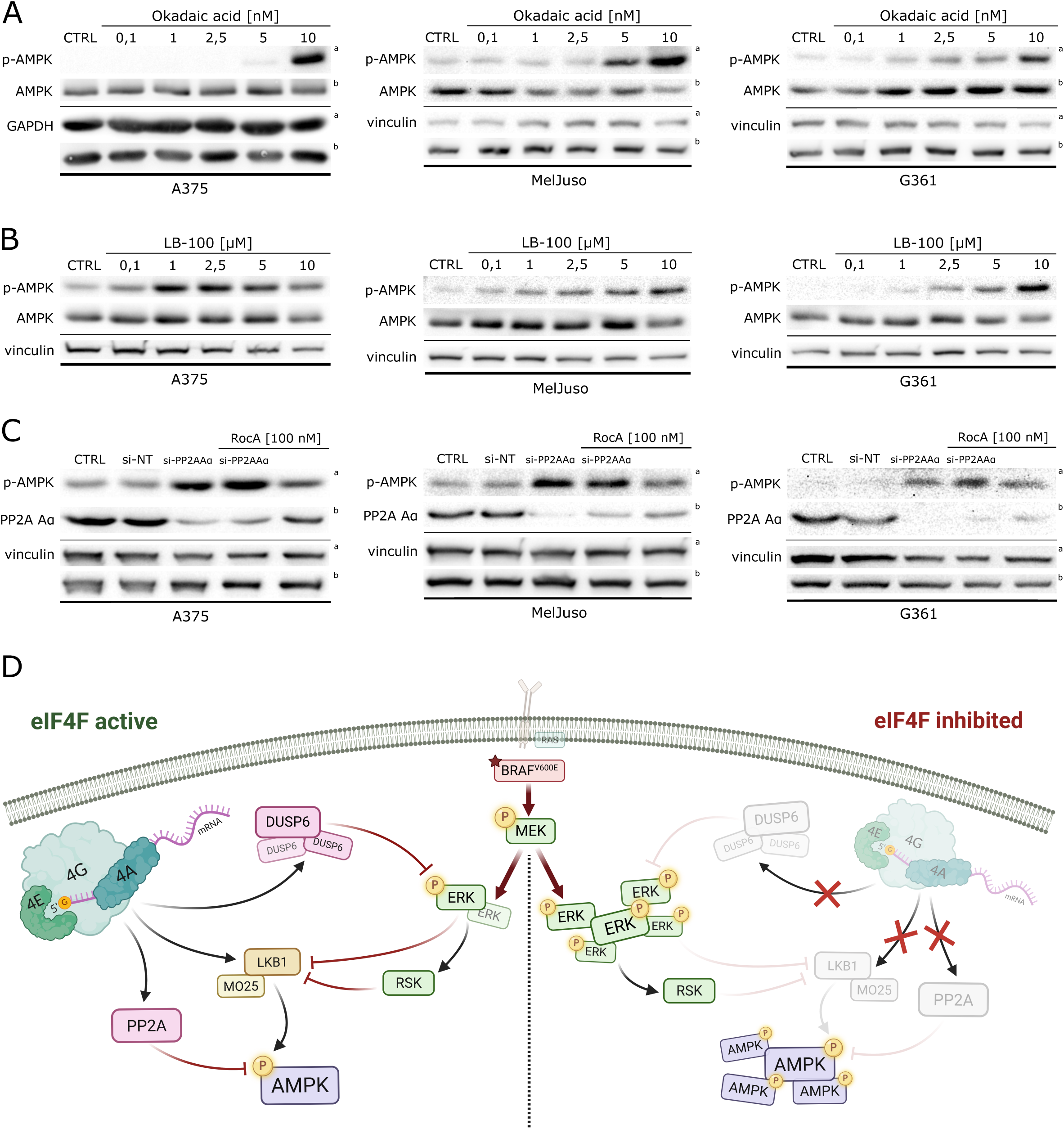
The eIF4F-PP2A-AMPK axis restricts AMPK activity in melanoma. Western blot analysis of A375, MelJuso, and G361 cells treated with increasing concentrations of the PP2A inhibitor **(A)** okadaic acid for 20 h and **(B)** LB-100 for 20 h. PP2A inhibition induced dose-dependent AMPK activation in all three melanoma cell lines regardless of LKB1 status. **(C)** Western blot analysis of A375, MelJuso, and G361 cells transfected with PP2A-Aα subunit-targeting siRNA for 72 h, with or without 100 nM RocA treatment (20 h). siRNA-mediated knockdown of PP2A-Aα subunit increased AMPK phosphorylation to levels comparable with RocA treatment, confirming that the eIF4F-PP2A axis restricts AMPK activity in melanoma cells. Non-targeting siRNAs (si-NT) were used as a control. Vinculin served as a loading control. The upper index letters refer to the corresponding loading control detected on the same membrane. **(D)** eIF4F activity controls AMPK signaling in melanoma bearing *BRAF* and *NRAS* mutations. Schematic model illustrating eIF4F-dependent control of ERK and AMPK signaling. Inhibition of eIF4F disrupts parallel regulatory axes, leading to ERK activation through loss of DUSP6-mediated negative feedback and to AMPK activation due to reduced PP2A-dependent negative regulation, independently of LKB1. Created with BioRender.com.

Taken together, these results provide a new mechanistic insight into the anti-cancer activity of compounds targeting eIF4F and indicate a therapeutic potential for small molecule eIF4F and PP2A inhibitors to activate the growth-inhibitory AMPK signaling in melanoma regardless of the LKB1 status of cancer cells.

## Discussion

The management of metastatic melanoma has been difficult due to its inherent resistance to conventional chemotherapy and radiotherapy. Pharmacological BRAF and MEK inhibitors are highly effective for advanced melanomas carrying the common BRAF V600E mutation and have extended patient survival (54, 55). Nevertheless, most patients eventually develop resistance, resulting in disease relapse and progression. Recent developments in immunotherapy, particularly combinations of immune checkpoint inhibitors, have significantly improved treatment outcomes, boosting 5-year survival for advanced melanoma to around 50 %. However, a large proportion of patients treated with immunotherapy still experience disease progression (55, 56).

The eIF4F translation initiation complex has been identified as a key factor in the emergence of melanoma persister cells and the development of resistance to clinical BRAF and MEK inhibitors (36, 57–59). Interestingly, eIF4F has also been implicated in regulating tumor microenvironment and melanoma immune escape (33, 60). This makes it a potential target for strategies to enhance melanoma sensitivity to both BRAF/MEK-targeted therapy and immunotherapy. In our recent study (37), we provided molecular insights into how the eIF4F complex contributes to the negative regulation of the ERK MAPK signaling pathway. We showed that eIF4F sustains the ERK negative feedback in melanoma by maintaining DUSP6/MKP3 protein levels. Disrupting eIF4F leads to a rapid drop in DUSP6 expression and drives extreme ERK hyperactivation in melanoma cells (37).

In this work, we present a new eIF4F-PP2A-AMPK axis acting in parallel to the eIF4F-DUSP6-ERK axis controlling MAPK activity (Fig. 5D). Inhibiting eIF4F simultaneously unleashes ERK signaling (by disarming the DUSP6-mediated negative feedback) and elevates AMPK activity (by diminishing the negative control by the PP2A phosphatase) (51, 52). Such unanticipated convergence between translational control, growth signaling, and energy-sensing pathways seems paradoxical in BRAF V600E melanoma, where ERK/RSK phosphorylation of LKB1 was reported to suppress LKB1-mediated AMPK activation (31, 61). Yet two of our results reconcile the paradox. First, AMPK activation upon eIF4F inhibition occurs in LKB1-null G361 cells and persists in LKB1 wild-type cells after LKB1 knockdown, establishing an LKB1-independent mechanism. Second, the magnitude of AMPK activation is modest compared with that induced by combined glycolysis and mitochondrial complex I inhibition (2-deoxyglucose plus rotenone), a condition that strongly suppresses upstream ERK signaling in BRAF V600E cells via AMPK-dependent disruption of BRAF–KSR complexes (32, 38). Thus, eIF4F inhibition appears to elicit a ‘sub-threshold’ AMPK response that can co-exist with amplified ERK signaling rather than extinguishing it.

Our proteomic screen uncovered downregulation of the PP2A Aα subunit (PPP2R1A) and UHRF1 following eIF4F inhibition. PP2A can directly dephosphorylate Thr172 and down-modulate AMPK activity (51, 62–64). Moreover, UHRF1 was recently shown to act as a nuclear gate-keeper of AMPK by promoting AMPK–PP2A association; loss of UHRF1 elevates AMPK activity (53). Consistent with these mechanisms, both PP2A inhibitors (okadaic acid and LB-100) and genetic depletion of the PP2A Aα scaffold increased P-AMPK levels in melanoma cells. Together, these data support a model in which eIF4F maintains expression of PP2A components and cofactors (including UHRF1), thereby restraining basal AMPK signaling; when eIF4F is inhibited, the PP2A brake weakens and AMPK phosphorylation rises. Importantly, the selective PP2A inhibitor LB-100 has entered clinical testing, and PP2A inhibition can synergize with cytotoxic cancer therapy and immunotherapy (65–67). Our new data adds ‘metabolic rewiring’ via AMPK to the list of LB-100’s potential mechanisms of action in cancer.

Our insights could inform several therapeutic strategies. First, combining partial eIF4F inhibition with MAPK-pathway inhibitors could shift the AMPK/ERK balance towards cytostasis while minimizing ERK rebound. Second, using transient PP2A inhibition (e.g., LB-100) could elevate AMPK activity and suppress ERK-driven proliferation in BRAF V600E tumors, including those lacking *LKB1*. Although we focused on melanoma, we also observed loss of LKB1, MO25, and PP2A Aα, along with AMPK activation in a *BRAF*-mutant colorectal cancer model. This suggests that the eIF4F–PP2A–AMPK axis may operate in other *BRAF*-addicted tumors.

We did not pinpoint the upstream kinase responsible for AMPK activation during eIF4F inhibition. The fact that eIF4Fi-induced AMPK activation is taking place in LKB1-deficient cells excludes the canonical LKB1-dependent route. Moreover, we observed that eIF4F inhibition lowers MO25/CAB39 and LKB1 protein levels, changes that would be predicted to reduce, not increase, canonical AMPK activation (9–12). We further excluded AMPK activation by CaMKKβ (18, 19, 68) using siRNAs, and because the CaMKKβ protein itself is eIF4F-dependent and decreases upon rocaglate exposure. Our data also suggest that the lysosomal protein complex Ragulator does not serve as a platform for AMPK activation upon eIF4F inhibition (41), as neither acute lysosome perturbation with the PIKfyve inhibitor YM201636 nor LAMTOR1 knockdown blunted the AMPK response to eIF4F inhibition, arguing that the v-ATPase–Ragulator axis is dispensable here (25, 26, 41). Moreover, our data indicate that AMPK activation by metabolic stress might also be LKB1- and Ragulator-independent in *BRAF*-mutant melanoma. This is unexpected given the published data obtained in other cell types. For example, AMPK activation by the mitochondrial complex I inhibitor metformin was reported to depend on v-ATPase, AXIN, and Ragulator/LAMTOR1 in mouse hepatocytes and fibroblasts (44). On the other hand, our data might reflect the use of 2-deoxyglucose to induce low-level metabolic stress and its combination with complex I inhibitor to induce high metabolic stress in our experiments, as 2-deoxyglucose could elicit endoplasmic reticulum stress in melanoma cells and stimulate AMPK activity via CaMKKβ (18, 69).

The reason why our proteomic screen did not yield any obvious candidate for the AMPK kinase may be that, in response to eIF4F inhibition, its kinase activity or subcellular localization changes rather than cellular levels. Alternatively, the canonical LKB1-mediated control of AMPK might compete with the alternative AMPK-activating kinase. Future work using kinome-wide gene knock-out screens could help identify the kinase responsible for the observed AMPK activity in melanoma. Alternatively, proximity-labeling phosphoproteomics could pinpoint changes in AMPK-interacting kinases and phosphatases after eIF4F blockade.

In summary, our study provides new mechanistic insights into the function of eIF4F-mediated translation in malignant melanoma. These findings could have implications for melanoma resistance to current therapies. We identify translational control as a key regulator of melanoma signaling homeostasis: eIF4F preserves ERK negative feedback via DUSP6 and maintains a PP2A-centered negative control over AMPK, while also sustaining the expression of its canonical activators. As a result, disabling eIF4F reveals latent ERK signaling capacity while simultaneously freeing AMPK from both canonical regulation and phosphatase control. Our findings could be therapeutically harnessed to restrain melanoma proliferation regardless of *LKB1* status.

## Materials and Methods

### Cell Culture and Treatments

Human melanoma cell lines A375 and G361 were purchased from the European Collection of Animal Cell Cultures (ECACC; Salisbury, UK), MelJuso and COLO800 were purchased from DSMZ. All cell lines were maintained at 37 °C in a humidified atmosphere containing 5% CO_2_ and cultured in RPMI 1640 medium (Sigma-Aldrich) supplemented with 10% fetal bovine serum (Sigma-Aldrich), 1% penicillin/streptomycin (Gibco), and 2 mM L-glutamine (Gibco). Cells were regularly tested for mycoplasma contamination using Mycoplasma Detection Kit (Biotool, Munich, Germany) and DAPI staining of fixed cells followed by immunofluorescence microscopy.

Small-molecule inhibitor stocks of Rocaglamide A, 4E1RCat, CR-1-31-B, Rotenone (MedChemExpress), PD184352, YM201636 (Selleckchem) and Okadaic acid (Merck) were prepared in dimethylsulfoxide (DMSO, Merck). 2-deoxyglucose (MedChemExpress) was dissolved in RPMI media and LB-100 (ApexBio) was dissolved in sterile MilliQ H_2_O. Inhibitor stocks were diluted in pre-warmed RPMI media and added to the cells. Control samples were treated with the corresponding amount of the vehicle.

#### siRNA-mediated knockdown of gene expression

Cells were transfected using the Lipofectamine™ RNAiMAX Transfection Reagent (Thermo Fisher Scientific) following the manufacturer’s instructions. The gene-specific siRNAs used for knockdown: LKB1 siRNA (sc-35816), eIF4E siRNA (sc-35284), eIF4G siRNA (sc-35286), eIF4AI siRNA (sc-40554), LAMTOR1 siRNA (sc-96597), PP2A Aα siRNA (sc-44033), CaMKKβ siRNA (sc-38956), MLK3 siRNA (sc-35946) from Santa Cruz Biotechnology, STK11 (4392420, ID: s13579) from ThermoFisher Scientific, and MISSION® esiRNA PPP2R1A (EHU071351) from Merck. AllStars Negative Control siRNA (1027281) was purchased from QIAGEN.

### Western blot

Cells were lysed in 2× Laemli sample buffer, and proteins in total cell lysates were resolved by SDS-PAGE (10 or 12,5% acrylamide) using vertical electrophoresis unit SE250 (Hoefer) and transferred to PVDF membranes (Merck) in the Trans-Blot SD semi-dry transfer system (Bio-Rad). Membranes were blocked with 5% non-fat milk in Tris-buffered saline + 0.1% Tween 20 (TBST) for one hour at room temperature and incubated with primary antibodies overnight at 4 °C. The next day, membranes were washed 5 × 5 min in TBST and incubated with horseradish peroxidase-linked secondary antibodies for one hour at room temperature. After 5 × 5 min TBST washing, proteins were visualized with enhanced chemiluminescence (ECL) substrate (Thermo Fisher Scientific) in the G:BOX detection system (Syngene). PageRuler™ Prestained Protein Ladder 10 to 180 kDa (26616, Thermo Fisher Scientific) was used as a protein size marker. All antibodies were used according to the manufacturers’ recommendations.

Primary antibodies used for western blot: monoclonal rabbit anti-phospho-AMPK (2535), monoclonal rabbit anti-AMPK (2532), monoclonal rabbit anti-phospho-ERK1/2 (#4370), polyclonal rabbit anti-ERK1/2 (#9102), monoclonal rabbit anti-EGR1 (#4153), monoclonal rabbit anti-eIF4A (#2013), monoclonal rabbit anti-eIF4E, polyclonal rabbit anti-eIF4G (#2498), monoclonal mouse anti-CDK1 (9116), monoclonal rabbit anti-CDK2 (2546), monoclonal rabbit anti-LAMTOR1 (8975), monoclonal rabbit anti-LKB1 (3047), monoclonal rabbit anti-MO25 (2716), monoclonal rabbit anti-MLK3 (2817), monoclonal rabbit anti-PP2AAsubunit (2041), monoclonal rabbit anti-TYMS (5449) purchased from Cell Signaling Technologies and monoclonal mouse anti-DUSP6 (sc-377070), monoclonal mouse anti-vinculin (sc-73614), monoclonal mouse anti-CAMKKβ (sc-271674), monoclonal mouse anti-GAPDH (sc-47724) purchased from Santa Cruz Biotechnology. Mouse anti-PCNA (PC-10) was kindly provided by Dr. Bořivoj Vojtěšek (MMCI Brno).

Secondary antibodies mouse anti-rabbit IgG-HRP (sc-2357) and mouse IgG kappa binding protein m-IgGκ BP-HRP (sc-516102, Santa Cruz Biotechnology) were used for the detection.

### Tumor tissue lysate preparation

Tumor samples were obtained in an *in vivo* experiment described previously (37). Briefly, male NOD.Cg-Rag1^tm1Mom^ Il2rg^tm1Wjl^/SzJ (NRG, The Jackson Laboratory, Bar Harbor, ME) mice aged six to eight weeks were anesthetized and injected intradermally with 5 × 10^4^ A375 cells (stably transfected with the ERK activity luciferase reporter construct pKROX24(MapErk)Luc) in 50 μl PBS. After 19 days, mice were treated with the eIF4F inhibitor CR-1-31-B (0.2 and 0.4 mg/kg, in sesame oil, i.p.). Control animals were treated with the vehicle. Tumors were collected after the second live-cell imaging, performed 24 h post-treatment. The experiments were conducted with the approval of the Academy of Sciences of the Czech Republic (AVCR 55/2024), overseen by the local ethical committee, and performed by certified individuals (R.V., O.V., K.So.).

Tumors were processed using the gentleMACS Dissociator system (Miltenyi Biotec) according to the manufacturer’s protocol for protein extraction from mouse tissues. Each tumor was first weighed, and ice-cold lysis buffer (1% SDS supplemented with protease and phosphatase inhibitors) was added in a ratio of 1 ml per 100 mg of tissue. Samples were transferred into C-tubes, briefly inverted to submerge the tissue, and homogenized using the gentleMACS Protein_01 program. The homogenates were centrifuged at 4000 × g for 5 min at 4 °C, and the cleared supernatants were transferred into clean microcentrifuge tubes. Lysates were stored at-20 °C until further processing for western blot analysis.

### Samples preparation for LC-MS analyses

The A375 and MelJuso cell lines were seeded in 10-mm dishes and, on the following day, treated with 100nM RocA, 500 nM PD184352/CI-4040 (MEKi), and their combination (DRG_MEKi). Negative controls (CTR) were treated with an equivalent volume of DMSO. 20 h post-treatment, cells were washed, collected, and pelleted. Individual cell pellets were solubilized using SDT buffer (4% sodium dodecyl sulphate, 100mM dithiothreitol, 100mM TrisHCl, pH 7.6) and processed by filter-aided sample preparation (FASP) method (70) with some modifications. The samples were mixed with 8M UA buffer (8M urea in 100 mM Tris-HCl, pH 8.5), loaded onto the Microcon device with MWCO 30 kDa (Merck Millipore), and centrifuged at 7,000× *g* for 30 min at 20°C. The retained proteins were washed (all centrifugation steps after sample loading done at 14,000× *g*) with 200 μL UA buffer. The final protein concentrates kept in the Microcon device were mixed with 100 μL of UA buffer containing 50 mM iodoacetamide and incubated in the dark for 20 min. After the next centrifugation step, the samples were washed three times with 100 μL UA buffer and three times with 100 μL of 50 mM NaHCO_3_. Trypsin (enzyme:protein ratio of 1:50, sequencing grade, Promega) was added onto the filter, and the mixture was incubated for 18 h at 37°C. The tryptic peptides were eluted by centrifugation, followed by two additional elutions with 50 μL of 50 mM NaHCO_3_. Peptides were then cleaned by liquid-liquid extraction (3 iterations) using water-saturated ethyl acetate (71).

1/10 of the cleaned FASP eluate was collected for direct LC-MS analysis and completely evaporated in a SpeedVac concentrator (Thermo Fisher Scientific). Peptides were further transferred into LC-MS vials using 50 μL of 2.5% FA in 50% ACN and 100 μL of pure ACN, and with the addition of polyethylene glycol (final concentration 0.001%), and concentrated in a SpeedVac concentrator.

Phosphopeptides were enriched from the remaining 9/10 of the cleaned FASP eluate after complete solvent evaporation (SpeedVac concentrator) using High-Select™ TiO_2_ Phosphopeptide Enrichment Kit (Thermo Fisher Scientific, Waltham, Massachusetts, USA) according to the manufacturer’s protocol. Resulting phosphopeptides were extracted into LC-MS vials by 2.5% formic acid (FA) in 50% acetonitrile (ACN) and 100% ACN with the addition of polyethylene glycol (final concentration 0.001%) and concentrated in a SpeedVac concentrator.

### LC-MS analysis of peptides

LC-MS/MS analyses of all peptide mixtures (2 peptide solutions for each sample – 1 not enriched; 2 enriched on phosphopeptides using TiO_2_ enrichment kit) were done using an Ultimate 3000 RSLCnano system connected to Orbitrap Fusion Lumos Tribrid mass spectrometer (Thermo Fisher Scientific). Prior to LC separation, tryptic digests were online concentrated and desalted using a trapping column (100 μm × 30 mm, column compartment temperature of 40°C) filled with 3.5-μm X-Bridge BEH 130 C18 sorbent (Waters). After washing of trapping column with 0.1% FA, the peptides were eluted (flow rate-300 nl/min) from the trapping column onto an analytical column (Acclaim Pepmap100 C18, 3 µm particles, 75 μm × 500 mm; column compartment temperature of 40°C, Thermo Fisher Scientific) by using 100 min long nonlinear gradient program (1-56% of mobile phase B; mobile phase A: 0.1% FA in water; mobile phase B: 0.1% FA in 80% ACN; 0 min: 1% B, 70 min: 30% B, 100 min: 56% B) for analysis of phosphopeptides enriched fraction and not enriched peptide mixture. Equilibration of the trapping column and the analytical column was done prior to sample injection into the sample loop. The analytical column outlet was directly connected to the Digital PicoView 550 (New Objective) ion source with the sheath gas option and a SilicaTip emitter (New Objective; FS360-20-15-N-20-C12) utilization. ABIRD (Active Background Ion Reduction Device, ESI Source Solutions) was installed.

MS data were acquired in a data-dependent strategy with a cycle time of 3 seconds and with a survey scan (350-2000 m/z). The resolution of the survey scan was 60000 (200 m/z) with a target value of 4×10^5^ ions and a maximum injection time of 50 ms. HCD MS/MS (30% relative fragmentation energy, normal mass range) spectra were acquired with a target value of 5.0×10^4^. The MS/MS spectra were recorded in Orbitrap at a resolving power of 30 000 (200 m/z), and the maximum injection time for MS/MS was 500 or 54 ms for analysis of a phosphopeptide-enriched fraction or non-enriched peptide mixture, respectively. Dynamic exclusion was enabled for 60 s after one MS/MS spectra acquisition. The isolation window for MS/MS fragmentation was set to 1.6 m/z. Inspect the raw data as part of the PRIDE upload for more details on methods used.

The analysis of the mass spectrometric RAW data files was carried out with MaxQuant software (version 1.6.1.0) using default settings unless otherwise noted. MS/MS ion searches were done against a modified cRAP database (based on http://www.thegpm.org/crap, 112 protein sequences) containing protein contaminants like keratin, trypsin, etc., and UniProtKB protein database for *Homo sapiens* (https://ftp.uniprot.org/pub/databases/uniprot/current_release/knowledgebase/reference_proteomes/Eukaryota/UP000005640/UP000005640_9606.fasta.gz; version 2018-05 from 2018-05-28, number of protein sequences: 20,996). Oxidation of methionine, deamidation (N, Q), acetylation (protein N-terminus), and phosphorylation (S, T, Y; only data for solutions enriched on phosphopeptides) as optional modifications, and trypsin/P enzyme with 2 allowed miss cleavages and minimal peptide length 6 aminoacids were set. Peptides and proteins with an FDR threshold <0.01 and proteins having at least one unique or razor peptide were considered only. The match between runs was set separately for all enriched and non-enriched peptide solution analyses.

Protein intensities reported in proteinGroups.txt file and evidence intensities reported in evidence.txt file (output of MaxQuant program) were further processed using the software container environment (https://github.com/OmicsWorkflows), version 3.6.1a. Processing workflow covered: 1) protein level data processing: a) removal of decoy hits and contaminant protein groups, b) protein group intensities log_2_ transformation, c) LoessF normalization of protein Intensity values and d) differential expression using LIMMA statistical test (qualitative changes were considered separately without statistical evaluation); 2) phosphopeptide level: a) removal of not phosphorylated and contaminant proteins associated evidences, b) grouping of the evidences for the identical combination of peptide sequence and set of modifications (grouped evidences intensity summed), c) grouped evidences intensities log_2_ transformation, d) LoessF normalization of transformed intensities, and d) differential expression using LIMMA statistical test (qualitative changes were considered separately without statistical evaluation). Phosphopeptide level data evaluation results and data obtained from MEKi and DRG_MEKi samples were not used within this publication.

Commonly downregulated proteins in response to 100nM RocA were subjected to functional enrichment and clustering analysis using Metascape (https://metascape.org) (50). The list of 57 proteins identified in both melanoma subtypes was assessed across Gene Ontology Biological Process and curated pathway databases, including Reactome and KEGG. Enrichment significance was evaluated using false discovery rate (FDR) correction as implemented in Metascape. To identify higher-order biological themes, enriched terms were clustered based on shared gene membership, and functionally related terms were grouped into broader biological modules for interpretation.

### 5-ethynyl-2’-deoxyuridine (EdU) flow cytometry cell proliferation assay

Cell proliferation was evaluated by measuring DNA synthesis using the Click-iT Edu Alexa Fluor 488 flow cytometry kit (Thermo Fisher Scientific). Cells were seeded on a 12-well plate and allowed recovering overnight. Next day, cells were treated with increasing concentrations of Rocaglamide A (10, 25, 50, 100 nM). Controls were treated with equivalent amount of the vehicle (DMSO). After 18 hours, the inhibitor-containing media was replaced with fresh RPMI allowing cells to recover for 90 minutes. The cells were then pulse-labeled with 10 µM EdU for 30 minutes. Non-labeled negative control was included. After incubation, the cells were washed, trypsinized and samples were processed according to the manufacturer’s instructions. Samples were immediately measured in a flow cytometer using a 488 nm excitation laser and 525/40 emission filter (CytoFLEX V2-B5-R3, Beckman Coulter) Three independent repetitions were measured and normalized to total protein concentration determined by Bio-Rad Protein Assay Dye Reagent used according to the manufacturer’s instructions (Bio-Rad). The data are presented as an average of EdU-positive population ± SE, p < 0.05.

## Acknowledgments

We thank Gabriela Filipová for technical assistance. We acknowledge CEITEC Proteomics Core Facility of CIISB, Instruct-CZ Centre, supported by MEYS CR (LM2023042, CZ.02.01.01/00/23_015/0008175, e-INFRA CZ (ID:90254)). This work was funded by the Czech Science Foundation (GA22-30397S, GA25-17766S) and the European Union – Next Generation EU-the project National Institute for Cancer Research (Programme EXCELES, Project No. LX22NPO5102).

## Author contributions

N.V., A.L., K.K., K.Sm., B.V., and F.K. designed and performed the experiments and analyzed the data. N.V. prepared the figures, figure legends, and materials and methods section. B.V., D.P., and Z.Z. performed proteomic analyses and analyzed the proteomic data. R.V., O.V., and K.So. handled animals and performed the *in vivo* experiment. S.U. conceived the study, designed the experiments, analyzed the results, and wrote the manuscript.

## Supplementary Data

**Supplementary Dataset S1.** Proteomic analysis of MelJuso (MJ) and A375 melanoma cells treated for 20 h with 100 nM Rocaglamide A (DRG), 500 nM PD184352/CI-4040 (MEKi), and their combination (DRG_MEKi). Negative controls (CTR) were treated with an equivalent volume of DMSO. (Provided as a separate MS Excel file)

**Supplementary Dataset S2.** Metascape analysis of proteins downregulated in response to RocA in both A375 and MelJuso cells. Functional enrichment and clustering analysis of 57 proteins commonly downregulated by RocA treatment in both melanoma subtypes. Analysis reveals convergence across core biological modules, including cell cycle regulation and DNA damage response; growth factor- and stress-associated signaling; cytoskeletal organization and cell adhesion; and metabolic and nutrient-responsive processes. The list of downregulated proteins included cyclin-dependent kinases (CDK1, CDK2, CDK6), proteins involved in DNA synthesis (PCNA, TYMS), and metabolic enzymes (ACLY, FASN), indicating coordinated suppression of proliferation-associated metabolic and cell-cycle-dependent pathways. (Provided as a separate MS Excel file)

